# Inverting proteomics analysis provides powerful insight into the peptide/protein conundrum

**DOI:** 10.1101/023515

**Authors:** Wilson Wen Bin Goh, Limsoon Wong

## Abstract

In proteomics, a large proportion of mass spectrometry (MS) data is ignored due to the lack of, or insufficient statistical evidence for mappable peptides. In reality, only a small fraction of features are expected to be differentially relevant anyway. Mapping spectra to peptides and subsequently, proteins, produces uncertainty at several levels. We propose it is better to analyze proteomic profiling data directly at MS level, and then relate these features to peptides/proteins. In a renal cancer data comprising 12 normal and 12 cancer subjects, we demonstrate that a simple rule-based binning approach can give rise to informative features. We note that the peptides associated with significant spectral bins gave rise to better class separation than the corresponding proteins, suggesting a loss of signal in the peptide-to-protein transition. Additionally, the binning approach sharpens focus on relevant protein splice forms rather than just canonical sequences. Taken together, the inverted raw spectra analysis paradigm, which is realised by the MZ-Bin method described in this article, provides new possibilities and insights, in how MS-data can be interpreted.

## Introduction

Mass Spectrometry (MS)-based proteomics is a critical technology in high-throughput biological and, more recently, clinical research (1). Applications include biomarker identification, drug-target identification, patient subtyping, and biological genotyping (2–4).

The basic principle of MS-based proteomics lies in the concurrent measurement of the mass-to-charge ratio (m/z) of large numbers of ionized peptides or peptide fragments. These raw spectral data are then mapped to in silico libraries of known peptides in order to make peptide identifications. Contemporary proteomics setups normally generate two sets of spectral data, MS1 and MS2. MS1 contains direct information (including the m/z) on ionized peptides. MS2 contains corresponding information on the peptide fragments, following molecular dissociation.

In a typical MS experiment, only about 10-30% of spectra are mappable to known peptides with sufficient statistical confidence (5). Of the remaining 70-90%, aside from noise, these theoretically contain data from mappable peptides that do not meet statistical thresholds, unknown peptides (which are not represented in the library) and post-translational modifications (PTMS) (which are difficult to reliably characterize).

Although it is a standard practice, mapping the experimental MS space to identify and quantify known proteins, if only to compare differences between sample classes (e.g. normal and cancer), is highly inefficient and error prone: Assignment of spectra to associated peptides (peptide-spectral matching/PSMs) usually requires a high level of statistical stringency due to the large number of erroneous matches. If the reference library is large, the statistical stringency threshold needs to be adjusted accordingly; this makes it harder to obtain statistically significant matches, which in turn reduces proteome coverage. In the event that PTMs and other modifications are considered, this increases the size of the search space (since both modified and unmodified peptide forms have to be included), as well as the complexity and error rate (e.g. the modification sites could include a wide range of amino acids with unknown probabilistic bias). Aside from size, different reference libraries (6) and different search algorithms (7) can give rise to different peptide/protein identification results.

Errors can arise in several situations. As mentioned, inclusion of PTMs, which requires increasing the tolerance range for match inclusion, decreases the reliability of matches. Another significant error source arises from stochastic selection of different peptides in different samples. The top n peptides are typically used to generate the final protein expression in a sample, but the constituent peptides in each sample may be different; this may lead to inaccurate quantitation (signal dilution) for the same protein amongst different samples. Moreover, it is naïve to consider canonical protein sequences wholesale (all exons present) without consideration of the possibility of splice variants (some exons are excluded).

Within a comparative experimental framework, we propose that only a minority of spectral features are expected to be strongly differential between the classes anyway (e.g. normal and cancer class). These spectral features alone may be sufficient for class prediction without peptide/protein matching. Differential spectral features are expected to be associated with biologically relevant features (e.g. peptides/proteins). Thus, identifying these differential spectral features first can theoretically reduce the spectral search space, thus leading to more efficient, reliable and focused identification of only relevant biological features (peptide and their corresponding proteins).

On the other hand, raw spectra is ridden with high redundancy rates and noise. These issues make it difficult theoretically to directly identify useful spectral features from raw spectra. Saeed et al have proposed a graph-theory based approach of identifying informative low-signal spectra “clusters” which in turn can be directed to library search algorithms (5). However, comparison of each spectrum with every other spectrum makes the clustering problem computationally costly. While this can be mitigated by parallelization, such a high-quality computing set-up is not feasible for most proteomic investigators (8).

The first aim of this work is to propose and demonstrate that inverting the conventional proteomics analysis strategy can be efficiently achieved (i.e., instead of “raw spectra-> PSMs->proteins->feature identification”, we do “raw spectra-> feature identification->PSMs + optimized de novo start sites”) via a simple m/z binning strategy. We term this approach MZ-Bin.

Recent technological advancements in proteomics have led to a surge of Data-Independent Acquisition (DIA) methods [19, 20]. These methods fragment precursors and record fragment ions independent of their stoichiometry in the sample, thereby offering more consistent protein quantification across samples. The different DIA methods include MSE (9) and SWATH (10). While the data-acquisition approach of these methods is conceptually similar, they differ in the way the data are analyzed. SWATH acquires data by repeatedly cycling through precursor isolation windows, referred to as SWATH windows, within a predefined m/z range covering the whole mass range of most MS-measurable precursors (10). The SWATH strategy thus creates an exhaustive multi-windowed SWATH map for every sample via a single injection. SWATH, as an instance of next-generation high-throughput proteomics, is beginning to see widespread usage both in clinical proteomics (11), dynamic pathway characterization (12) and human proteome characterization (13). Thus, we demonstrate MZ-Bin’s performance on large complex data generated by SWATH-MS, as well as explore aspects of its idiosyncracies.

The second aim of this work is to demonstrate how MZ-Bin can produce interesting insights on the peptide/protein conundrum. Protein quantification is a key aspect of proteomic profiling yet this key goal is achieved via largely patchy information on constituent peptides. Protein splice forms are largely ignored in proteomic profiling. We wish to understand what is the cost, in terms of analytical reliability, of this naïve approach.

## Materials and Methods

### SWATH data

The SWATH dataset from (14) was used in this study. This dataset contains 24 SWATH runs from 6 pairs of non-tumorous and tumorous clear-cell renal carcinoma (ccRCC) tissues, which have been swathed in duplicates (12 normal, 12 cancer).

A control dataset (normal class only) is used for false-positive analysis. This is composed of 12 SWATH runs from a human kidney test tissue digested in quadruplicates and each digest analyzed in triplicates using a tripleTOF 5600 mass spectrometer (AB Sciex).

### SWATH data interpretation

All SWATH maps for the renal cancer dataset were analyzed using OpenSWATH (13).

A spectral reference library containing 49,959 reference spectra for 41,542 proteotypic peptides was generated from 4,624 reviewed SwissProt proteins (14). The spectral library was compiled using DDA data of the kidney tissues in the same mass spectrometer.

The peptides identified were aligned prior to protein inference using the algorithm TRansition of Identification Confidence (TRIC) (version r238), which is available from https://pypi.python.org/pypi/msproteomicstools and https://code.google.com/p/msproteomicstools. The parameters used for the feature_alignment.py program are: max_rt_diff=30, method=global_best_overall, nr_high_conf_exp=2, target_fdr=0.001, use_score_filter=1.

As a standard practice in the SWATH pipeline, the signal intensity for the two most abundant peptides per sample were used to quantify proteins. 3,123 proteins were quantified across all samples with peptide and protein FDR below 1%.

### MZ-Bin strategy

#### 1/ Binning procedure

For each sample’s mz file, the individual m/z reads are binned and their corresponding intensities summed. The binning process is performed in tiers (level 1….level n).

At level n, an m/z value x is mapped to the MZ-bin floor((10^(n-1)^ * x)+0.5)/10(n-1). E.g., at level 2, the values 10.1, 10.12, 10.123,… are all mapped to the MZ-bin 10.1 and, at level 3, the values 10.12, 10.123, 10.1234,… are all mapped to the MZ-bin 10.12.

Each MZ-bin contains information on the summed intensities for all m/z reads in that MZ-bin.

#### 2/ Feature selection

To improve the feature-selection process, a three-step rule-based feature-selection procedure was designed. In the first step, if an MZ-bin has non-zero intensity values in more than half of the tissues in class A and non-zero in more than half of the tissues in class B, it is kept. Otherwise, the MZ-bin is discarded.

In the second step, for each class of tissues, the top 20% MZ-bins supported by at least half of the tissues in that class are selected. That is, there is a subset of tissues that constitute more than half of the tissues in that class and, for each tissue in this subset, the MZ-bin’s summed intensity is among the top 20% of all MZ-bins in that tissue.

Only when the first two criteria have been met are significant MZ-bins selected, in the third step, based on the standard two-sample t-test (class A vs class B).

A t-statistic (*T*_*p*_) is calculated for each MZ-bin by comparing the summed intensity values between classes *C1* and *C2*, with the assumption of unequal variance between the two classes (15). *T*_*p*_ can be expressed as:

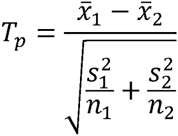

where 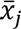 and *s_j_* are respectively the mean and standard deviation of the summed intensities of a given MZ-bin at some level n, and *n_j_* is the sample size, in class *C*_*j*_.

The *T*_*p*_ is compared against the nominal t-distribution to calculate the corresponding p-value. A feature is deemed significant if p-value ≤ 0.01 and thus selected into the next round of bin expansion.

The first filter ensures that only good-quality MZ-bins with few data holes are admitted for feature testing. The second filter considers only the strongest signals that are consistent across samples from the same class. Low-intensity peaks have more uncertainly and therefore, more likely to generate false positives. Moreover, since the data originates from clinical samples, inter-sample heterogeneity is expected to be an issue.

To demonstrate that the MZ-Bin feature-selection approach is useful, we also performed feature selection simply using the t-test alone (standalone t-test); i.e., without the first two rules.

#### 3/ Bin expansion

For the set of significant features selected at the (n-l)th level, we iteratively work up to the n-th level, and reselect the critical features based on the t-test at 99% significance level. For example, we can only select MZ-bin 10.123 provided we have earlier selected MZ-bin 10.12, and we can only select MZ-bin 10.12 provided we have even earlier selected MZ-bin 10.1, and we can only select MZ-bin 10.1 provided in the beginning we have selected MZ-bin 10.

Because the limit of resolution for m/z is up to four decimal places, the iterative MZ-Bin expansion procedure can theoretically proceed up to 4 levels (e.g. 10, 10.1, 10.12, 10.123).

#### 4/ Peptide inclusion

For each iteration of MZ-Bin (from Level 1 onwards), we can identify a set of peptides from the reference library that overlap with each significant MZ-bin based on their m/z.

### Cross-Validation (CV) accuracy evaluation

In CV, we split the data equally into 2 sets (training and testing; n=12 each) and performed feature selection using MZ-Bin. Each split was kept to the same proportion of cancer and normal tissues as in the original unsplit data set. We trained a Naïve Bayes classifier on the significant features identified from training set, and measured the accuracy of the trained classifier on the test set, where

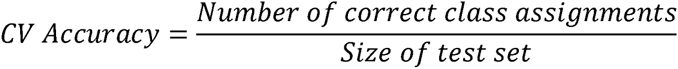

This was repeated 10 times on 10 different random splits of the data into training and testing sets.

To determine if the CV accuracy is meaningful, we randomly picked an equal number of features 1,000 times and retrained the Naïve Bayes classifier to produce a vector of null accuracy values. The CV Accuracy p-value is the number of times null accuracy ≥ significance threshold divided over total number of simulations.

The CV accuracy and the CV p-value can be combined by taking the ratio of the average CV accuracy and CV p-value. A method with high accuracy and low p-value thus generates a higher score.

## Results and discussions

### Implementation of the MZ-Bin strategy and theoretical underpinings

The principal concept underpinning MZ-Bin lies in the summarization of the mass spectra along the m/z dimension to generate MZ-bins at various levels of resolution (Level 1 to 4; see **Materials and Methods**) (Figure 1). Within each MZ-bin, the intensities of the mass ions are summed to generate an overall intensity for the MZ-bin. Differential MZ-bins are iteratively identified using some statistical comparison approach (in this paper, we used two approaches: a rule-based version and, for comparison, the standalone t-test followed by MZ-bin expansion). (See Figure 2 and Materials and Methods).

**Figure 1.**
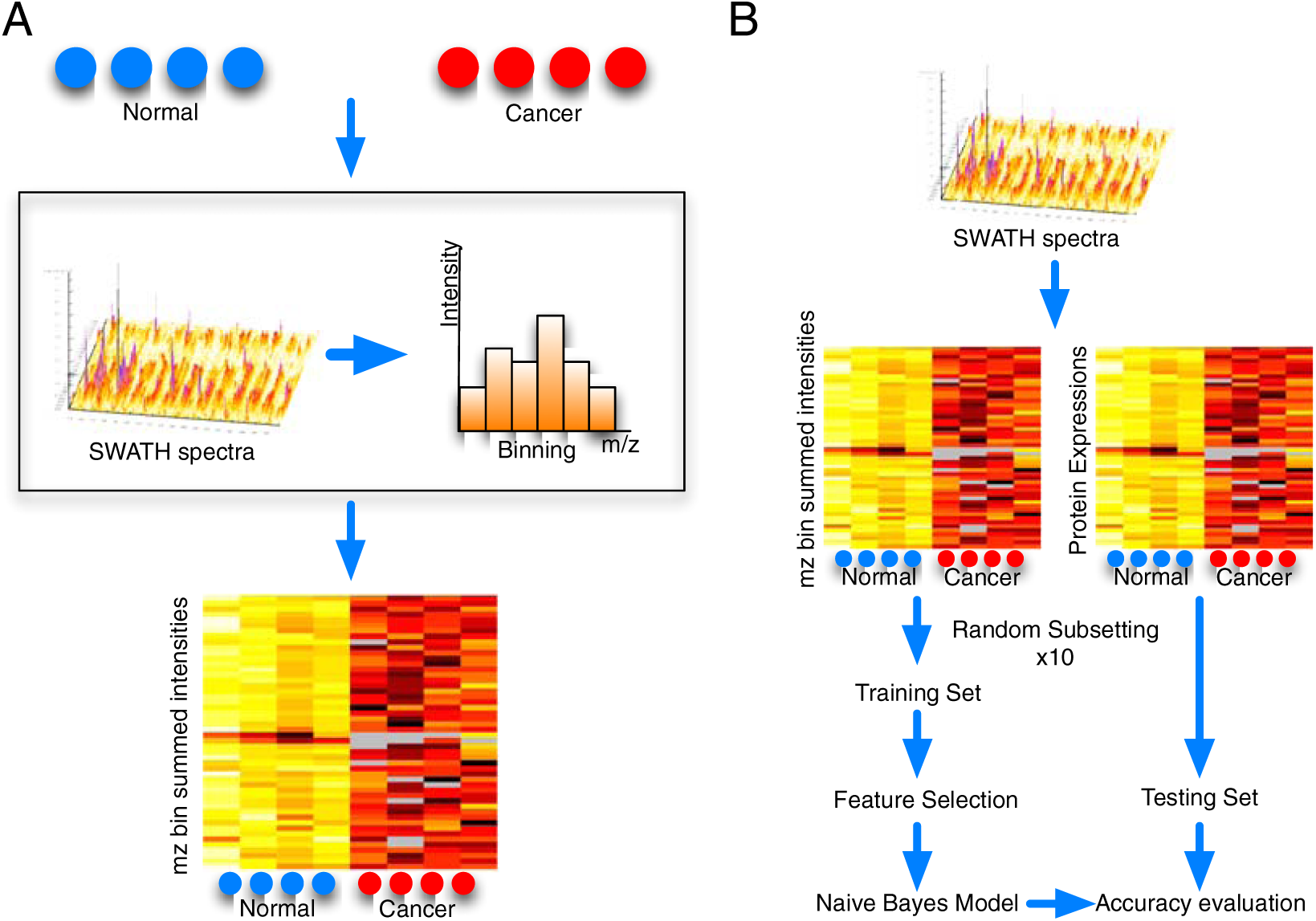
Proposed strategy for MZ-Bin A: Core Hypothesis. The MS profile can be compressed along the m/z dimension to generate informative MZ-bins that can discriminate between the sample classes (normal and cancer in this instance). These MZ-bins can be iteratively expanded to identify the relevant spectral features. **B: Naives Bayes Cross-Validation (CV) comparison between protein-based and spectral-based (MZ-Bins) feature selection.** To determine that the MZ-Bin strategy identifies relevant features, we compare this to conventional protein-identification strategy using CV accuracies generated across 10 random splits of the original dataset into training and testing sets.

**Figure 2.**
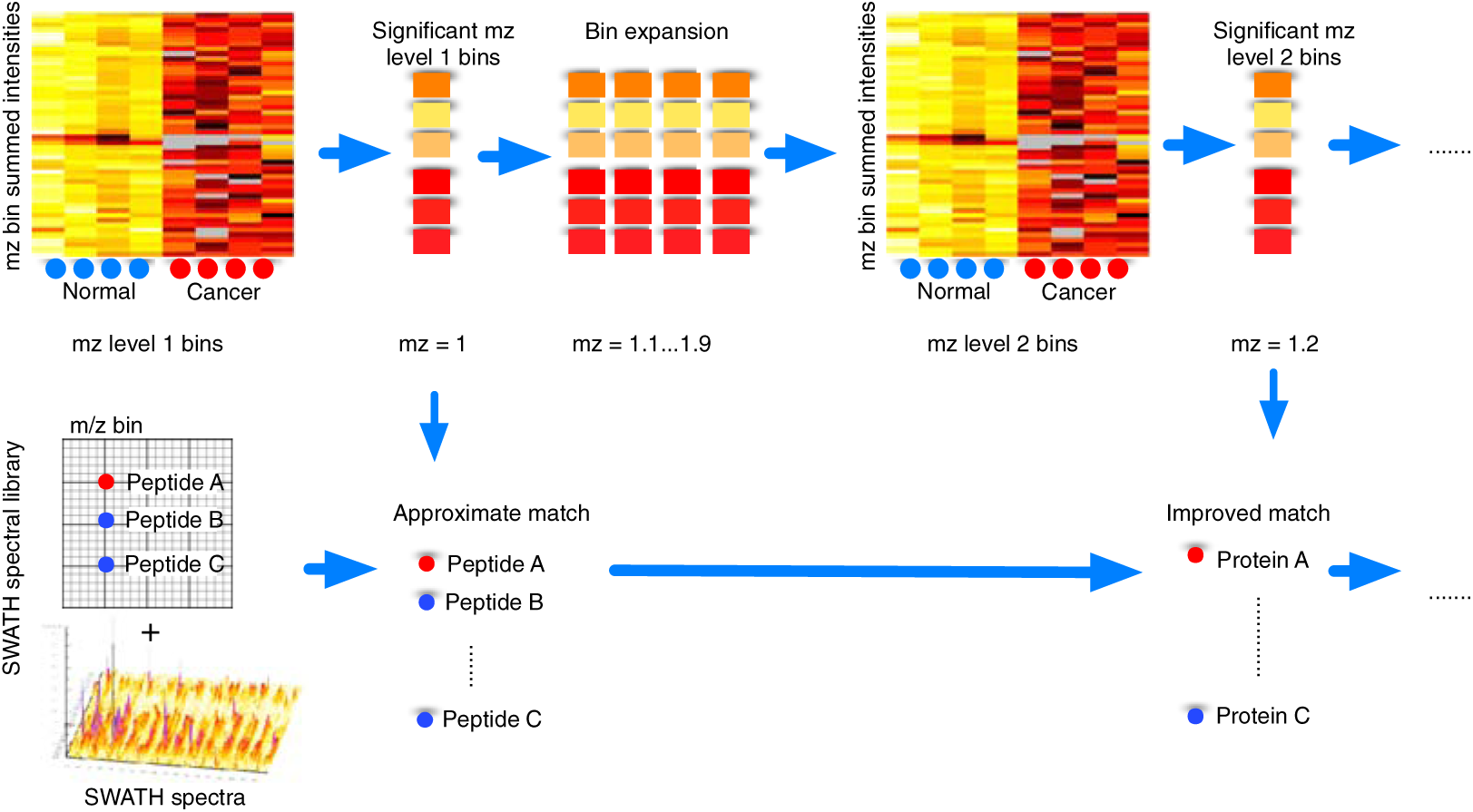
Iterative MZ-Bin expansion coupled to peptide matching. In our proposed strategy, the raw spectra are compressed along the m/z dimension, and the intensities summed per sample to generate the level-1 MZ-bins (integer). Significant MZ-bins are selected, and expanded into level-2 MZ-bins (1^st^ decimal), followed by recalculation of significant level-2 MZ-bins. This is repeated until level 4 (where we achieve an m/z range resolution of 3 decimal places). The iterative MZ-Bin expansion procedure should theoretically allow us to zoom into spectra corresponding to relevant peptides/proteins. To confirm this, we condensed the SWATH spectral library along the MZ dimension, and calculated the set of corresponding peptides/proteins corresponding to each set of significant MZ-bins (from level 1 to 4).

The relevance of the MZ-bins can be evaluated by identifying the peptides associated with the MZ-bins (mapping by m/z values). That is, we used the spectral library map generated by DDA (see SWATH data interpretation) that provides us with the expected m/z value and matched this with the significant MZ-bins.

Binning helps to resolve stochastic variation issues between runs. Performing feature selection without bins is difficult as there are many data holes due to stochastic variation and/or misalignments (Supplementary Figure 1A). Moreover, when the signals from each feature are not aligned properly, these features become uninformative, leading to poor cross-validation accuracy (Supplementary Figure 1B).

We hypothesize, based on the following assertions, that the intensity values encapsulated by these MZ-bins are informative:

1/ The m/z for a given peptide might shift within a range depending on the experimental, technical conditions, or due to stochastic variation. However, these shifts are likely to fall within the same MZ-bin (in particular, higher-resolution bins). Summing the m/z intensities in the bin can help recover from these measurement imperfections.
2/ We do not expect most proteins to be differentially expressed. Hence, even if peptides from different proteins X and Y fall into the same bin, in a comparison between classes A and B, it is often the case that only X is differentially expressed (say up-regulated in cancer) but Y is not, making the bin dominated by the peptide from X. Thus summing m/z intensities of X and Y helps wipe out the irrelevant Y.

To make a case for assertion 1, we considered the effects of feature selection without binning. Supplementary Figure 1A shows an example of the existence of data holes in imperfectly aligned spectral features. Moreover, without binning, the features are uninformative, and lack predictive power (Supplementary Fig 1B).

To make a case for assertion 2, the distribution of signal intensities for each m/z species within each level-1MZ-bin is shown in Supplementary Figure 1C. Here, level-1 MZ-bins from 400 to 1,200 m/z are shown with their constituent intensities sorted from highest to lowest. Intensities are z-normalized for cross-comparison purpose between level-1MZ-bins. For each MZ-bin, the signal intensity is overwhelmingly attributable to a very small number of m/z species, possibly belonging to the same peptide. The violinplot in Supplementary Figure 1C further shows that signal dominance is controlled by a small number of features --- e.g. on average, only 20 spectral features are needed to account for 25% of total intensity.

Thus, summing along the m/z dimension has the following benefits: 1/ Increase of signal for a given spectra location, 2/ elimination of redundancy, 3/ reduction of raw-spectra search space and consequently, computational power required for inter-sample class comparisons.

### Iterative bin expansion can only proceed up to level 3

Using the rule-based feature-selection approach, we monitored the number of spectral features filtered using rule 1 to 3, where rule 1 eliminates features that have data holes in more than half of the constituent samples of a class, rule 2 selects only the top 20% features supported by at least half in a class, and rule 3 is the two-sample t-test selection at p <= 0.01 (see **Materials and Methods**).

From level 1, we started with 1,200 MZ-bins. Rule 1 did not eliminate any features. Rule 2 reduced the feature set to 255. And we were left with 127 features post t-test.

This is expanded to form 1,143 features in level 2. Again, there were no obvious data holes, and rule 2 reduced the feature set to 414. From this, 183 features were selected post t-test.

In level 3, we started with 1,639 features (expanded from 183 level-2 MZ-bins), which were reduced to 414 features following rule 1 and 2. Interestingly, a large proportion of these (337) were significant. This suggested that we were increasing the enrichment for differential features as we worked downwards.

These 337 features can be expanded to 3,033 level-4 MZ-bins. Only at level 4 are a large number of features lost due to a large proportion of features having data holes exceeding more than half of a class. Only one MZ-bin passed the rule-1 check. This indicates that we hit the limit of resolution at level 3.

### Rule-based feature selection selects significantly fewer features yet maintains CV accuracy

To be more concise, we checked the effects on cross-validation accuracy for both the rule-based selection and standalone t-test selection for levels 2 to 3 (Table 1). Note that in SWATH, there are 32 m/z windows. The “merged” in the data tables means that we consider the m/z’s across all windows concurrently. Unsurprisingly, the number of features selected using rule-based selection is far lower than t-test alone. On average, at level 2, the rule-based method is selecting about less than half as many features selected using t-test alone. While in level 3, it is about seven-to-eight times lower. However, between levels 2 and 3, the accuracy of the resulting classifier is maintained, at about 80% for both rule-based and standalone t-test feature selection.

**Table 1.**
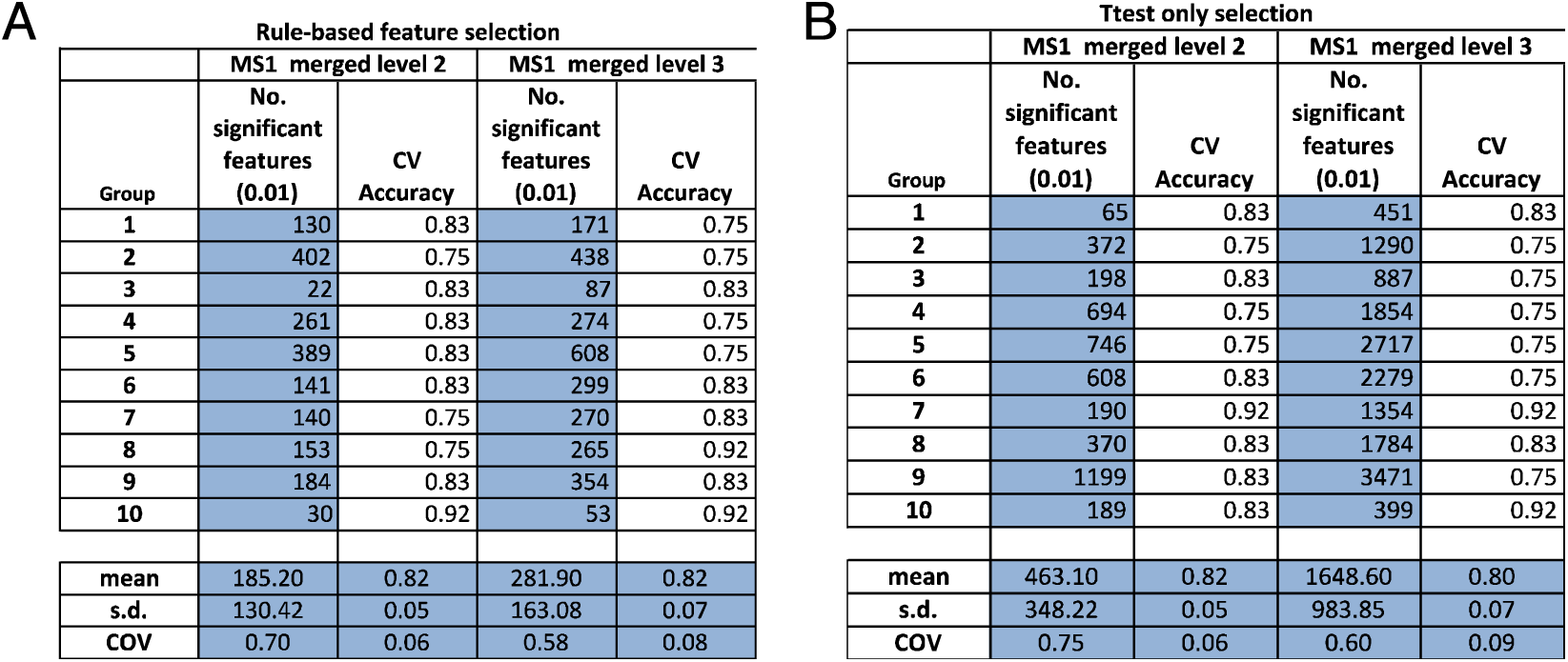
**Cross-Validation (CV) accuracies of level-2 and level-3 MZ-bins with and without rule-based feature selection**. The rule-based MZ-bin selection process expectedly generally predicts lower number of features than non-rule based (standalone t-test), while maintaining similar CV accuracy. This observation is consistent, as shown for levels 2 and 3.

### MZ-Bin false-positive rate is low

Using the normal class alone, we performed 1,000 random splits of the data and calculated the number of significant features identified at MZ-bin level 1 using the standard two-sample t-test (using the 95% significance level for illustration). These significant features are considered false positives since there is only one true class.

In this experiment, we considered the 32 SWATH windows concurrently, and included all m/z reads as long as they are within the m/z range for each MZ-bin. There are approximately 800 level-1 MZ-bins between 400 to 1200 m/z. Given these MZ-bins, the number of false positives is within expectation, with a median of 1 and mean of 34. These are within the expected number of false positives (expected value = 800 *0.05 =40). The standalone t-test is expected to be less stringent than our rule-based feature-selection method. Applying the rule-based feature-selection strategy reduced the number of false-positives further such that the median is now reduced to 0, and the mean is 0.14.

The experiment above confirms that the MZ-Bin approach does not generate a high number of false positives and that the rule-based strategy is highly stringent with very low false-positive rates (Figure 3A).

**Figure 3.**
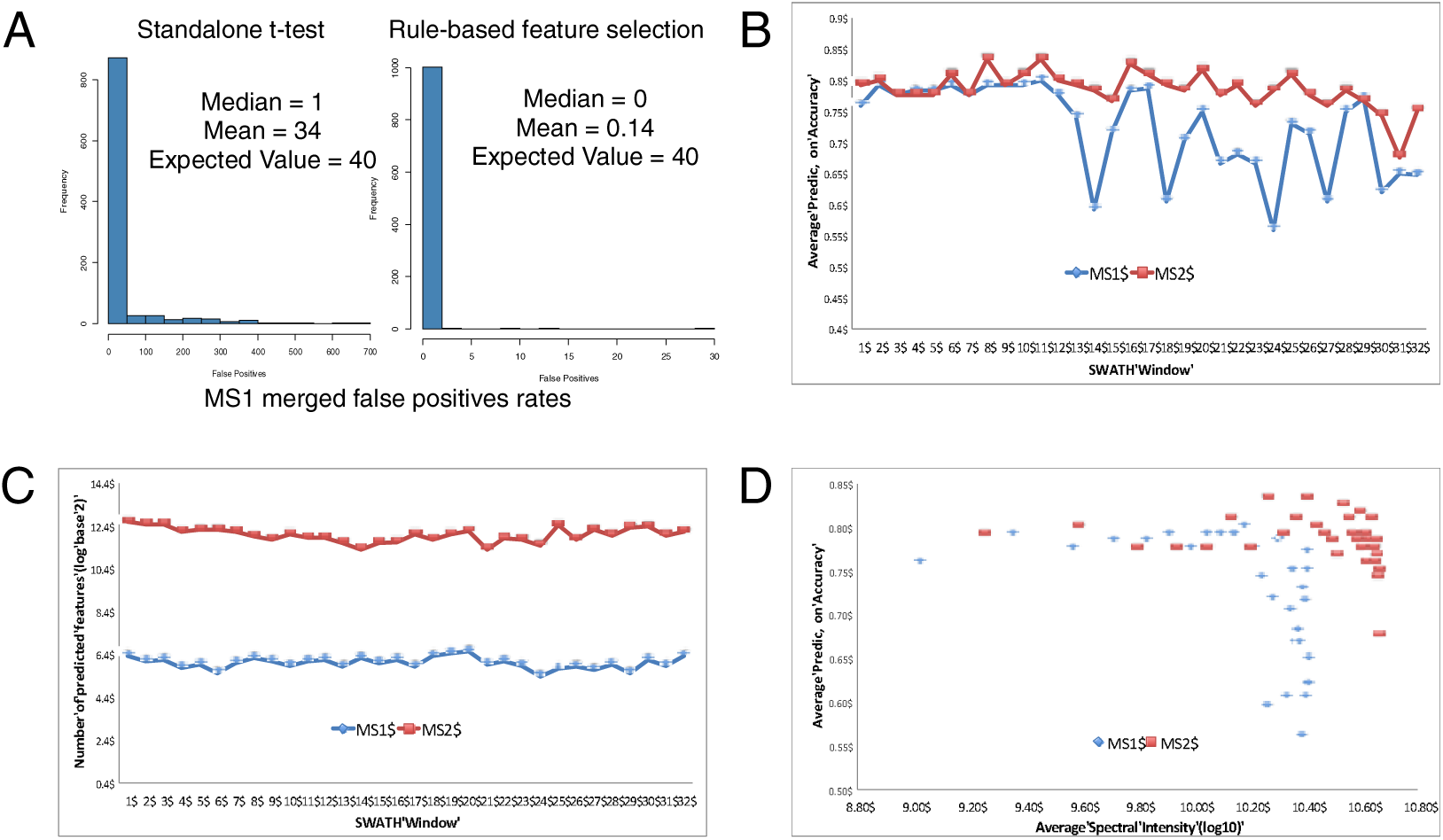
MZ-Bin false-positive rates and agreements of MZ-Bin performed on individual SWATH windows. A: False-positive checks. Using control data derived from normal tissue only, we randomly split these into two groups, and tested the level-1 MZ-bins 1,000 times using standalone t-test and rule-based feature selection. Note that the “merged” MZ-bins refers to concurrent analysis of the 32 SWATH windows. The number of false-positives is within expectation, with a median = 1 and mean = 34 (expected value = 800 *0.05 =40), confirming that the MZ-Bin approach does not generate overly high noise levels. However, the rule-based feature-selection strategy is even more stringent, with lower false positive rate**. B and C: Cross-validation and number of predicted features across 32 individual SWATH windows for both MS1 and MS2 level-1 MZ-bins**. MS1 are m/z values derived from unfragmented peptide species while MS2 are m/z values derived from fragmented peptides. We compared the MS1 and MS2 spectra over 32 SWATH windows and found that the patterns of cross-validation (CV) accuracy and number of predicted features are fairly similar demonstrating the consistency of the MZ-Bin strategy. **D: High-intensity SWATH windows have higher CV accuracy dispersal.** It is particularly interesting that SWATH windows with extremely high spectral intensities have wide CV accuracy range. This suggests that these SWATH windows are noisy and should be cleaned prior to feature selection.

### The feature identified and cross-validation (CV) accuracy are mirrored closely for MS1 and MS2

Although in this paper, we focus on MS1, the raw SWATH spectra contains both MS1 and MS2 data. As a standard rule-of-thumb in proteomics analysis, MS1 contains direct data on peptide ions while MS2 the corresponding fragmentation data, which is more complex and noisy. In addition, as an idiosyncrasy of SWATH, both MS1 and MS2 can be split into 32 windows based on m/z.

We checked that MS1 and MS2 average CV prediction accuracies and numbers of selected features by MZ-Bin across SWATH windows are mirrored closely (Figure 3B and 3C respectively). This confirms that the MZ-Bin approach generates consistent results irrespective of the SWATH window analyzed and between MS1 and MS2.

Using the standalone t-test on level-1 MZ-bins, we also considered whether there was substantial difference between the windows merged and unmerged (i.e., considering the SWATH windows concurrently or separately) in the cross-validation scenarios (Table 2). If the SWATH windows were perfectly discrete (and therefore non-overlapping), the merged and unmerged features should be similar. In practice, it is not so. Table 2 shows that across cross-validations, the merged windows generally select more features than unmerged while the cross-validation accuracy is maintained. This suggests that the windows are not discrete, and there are features that are only observable when the m/z’s are summed across all windows.

**Table 2.** **Cross-Validation (CV) accuracies of unmerged, unmerged SWATH windows, and protein-based expression (Single Proteins, SP)**. MS1 merged and unmerged level-1 MZ-Bins do not agree on the number of significant features, suggesting signal spillover between SWATH windows. Although the CV accuracy is high in SP, any random selection of random proteins yields equally high CV accuracy (cf. Supplementary Fig 2). On the other hand, although CV accuracy of MZ-Bin is lower, the results are statistically more meaningful.

| Group | MS1 unmerged level 1 spectra |  |  | MS1 merged level 1 spectra |  |  | Single Protein |  |  |
| --- | --- | --- | --- | --- | --- | --- | --- | --- | --- |
|  | No. significant features (0.05) | CV Accuracy | CV p-value | No. significant features (0.05) | CV Accuracy | CV p-value | No. significant features (0.05) | CV Accuracy | CV p-value |
| 1 | 95 | 0.83 | 0.00 | 103 | 0.83 | 0.00 | 901 | 1.00 | 0.92 |
| 2 | 178 | 0.83 | 0.00 | 228 | 0.83 | 0.00 | 1213 | 1.00 | 0.911 |
| 3 | 326 | 0.75 | 0.00 | 397 | 0.75 | 0.00 | 908 | 1.00 | 0.919 |
| 4 | 567 | 0.75 | 0.00 | 757 | 0.75 | 0.00 | 1210 | 1.00 | 0.905 |
| 5 | 302 | 0.75 | 0.00 | 376 | 0.75 | 0.00 | 800 | 1.00 | 0.915 |
| 6 | 203 | 0.83 | 0.00 | 237 | 0.83 | 0.00 | 1825 | 0.75 | 0.892 |
| 7 | 120 | 0.92 | 0.00 | 137 | 0.92 | 0.00 | 1017 | 1.00 | 0.902 |
| 8 | 208 | 0.83 | 0.00 | 248 | 0.83 | 0.00 | 1284 | 1.00 | 0.925 |
| 9 | 532 | 0.75 | 0.00 | 654 | 0.75 | 0.00 | 1251 | 1.00 | 0.917 |
| 10 | 245 | 0.83 | 0.00 | 321 | 0.83 | 0.00 | 834 | 1.00 | 0.92 |
| mean | 277.60 | 0.81 | 0.00 | 345.80 | 0.81 | 0.00 | 1124.30 | 0.98 | 0.91 |
| s.d. | 160.20 | 0.06 | 0.00 | 212.41 | 0.06 | 0.00 | 306.48 | 0.08 | 0.01 |
| COV | 0.58 | 0.07 | 0.00 | 0.61 | 0.07 | 0.00 | 0.27 | 0.08 | 0.01 |

To calculate a significance value for the observed cross-validation accuracy, we randomly picked a similar number of selected features 1,000 times and generated a null distribution of cross-validation accuracies. The p-value is thus the number of times the observed accuracy is lower than accuracy generated from randomly picked features divided over the number of simulations. We used the 99% significance level for this test. In no scenario did the randomly picked MZ-bins generate an accuracy greater than the observed (Supplementary Figure 2). Hence, the observed cross-validation accuracy is highly significant.

For comparison, a more conventional analysis of z-normalized protein expression using the two-sample standard t-test (Single Proteins, SP) was also performed. This conventional analysis generated many apparent false positives. On the full dataset, 53% of the 3,123 proteins were reported as significant by SP at the 99% significance level. For each of the cross-validations, a large number of protein features were also generated (Table 2). Although it seemed that the protein features generated very high cross-validation accuracy, almost any random subset of proteins (containing 5, 20 or 100 randomly picked proteins) generated strong CV accuracy when, in fact, these features were picked randomly and thus should be irrelevant to the classes. Generation of the null distributions for SP confirmed this (Supplementary Figure 2). In contrast, using MS1 merged level-1 MZ-Bins as illustration (where merged means we considered all SWATH windows concurrently), random selection of features did not generate null distributions with a strong left skew (i.e., very few randomly picked sets of MZ-bins generated null CV accuracies of 0.95 and above); cf. Supplementary Figure 2.

As a further control, we considered the effects of merged and unmerged windows in MS2 spectra (Supplementary Table 1). Since the MS2 m/z’s are generated from fragmented MS1 peptides, we expect much more redundancy. At the 99% significance level, the MS2 unmerged features averaged about 14,000 while merging reduced this to only about 600. But because these MS2 features are not easily mappable to any particular peptide (and highly redundant), they can only be used for pure profiling but not biological characterization. For this purpose, we used MS1-generated MZ-bins.

### High-intensity SWATH windows are the source of the non-discrete signals

Although MS1 and MS2 results are mirrored closely, some MS1 and MS2 windows have lower CV accuracies. We hypothesized that those windows with lower CV accuracies are associated with overall higher intensities. A plausible explanation for this disparity is that these windows may be enriched for MZ-bins that contain signal from more than one important contributing peptide, e.g. one peptide that is upregulated in disease, and the other in normal, which breaks assertion 2 (see above). Another possibility is that of spillage across MZ-bins, which is described below.

Figure 3D demonstrates that although average CV prediction accuracy is maintained across a wide range of average spectral intensities. The CV prediction accuracy becomes inconsistent and unreliable when average spectral intensity of the MZ-bins in a window is high. This is consistent for both MS1 and MS2.

Theoretically, SWATH captures m/z reads in discrete windows from the range of 400 to 1200 m/z. Therefore, if there are no signal overlaps between SWATH windows, then the number of significant features selected from merging all MS1 windows (MS1 merged), and maintaining the discrete windows (MS1 unmerged) should be similar.

However, in the earlier section we found that this is not so. Incorporation of all SWATH windows (including the high-intensity SWATH windows) generates different results in MS1 merged and MS1 unmerged, confirming signal overlaps between SWATH windows (Table 2).

As an arbitrary threshold, MS1 SWATH windows with average log10 spectral intensities above 10.35 and average CV prediction accuracies below 0.65 were removed (Figure 3D). Removal of these high-intensity but low CV accuracy windows allows MS1 merged and MS1 unmerged results to perfectly agree (Supplementary Table 2). This suggests that the lower intensity SWATH windows are perfectly discrete, with no signal spillage. Perhaps, as an idiosyncracy of the SWATH platform, to clean the SWATH windows in a less arbitrary manner, avoid losing whole windows (and their associated features), and double counting stray m/z intensities, we recommend clipping off all m/z values reported in any SWATH window that is outside its associated range.

### MZ-Bin associated peptides have stronger signal than their corresponding proteins

Across the 3 MZ-bin levels, we counted the number of associated peptide groups and corresponding proteins based on the reference DDA map (Figure 4A). In accordance with expectation, as the MZ-bin resolution increases from level 1 to level 3, the number of significant bins also increases. On the other hand, this leads to a concomitant decrease in the number of peptide groups, and corresponding proteins. This is not surprising because a lower-resolution MZ-bin (e.g., the level-1 MZ-bin 10) corresponds to a set of higher-resolution bins (e.g., the level-3 MZ-bins 9.50,…, 10.49). When MZ-bin 10 is significant, it might be due to several of the corresponding level-3 MZ-bins being significant, but not necessarily all of its corresponding level-3 MZ-bins. Thus while peptides whose m/z values are in MZ-bin 10 would be declared significant when the analysis was performed at level 1, they might not be declared significant when analysed at level 3 if their m/z values were not in any of the corresponding level-3 MZ-bins that are significant. In our experiment, the set of MZ-bins selected at level 3 mapped to 1,044 peptide groups and 688 corresponding proteins. This is much more specific than if the t-test were applied directly on the identified protein abundance levels where 1,649 out of 3,123 proteins were reported significant.

**Figure 4.**
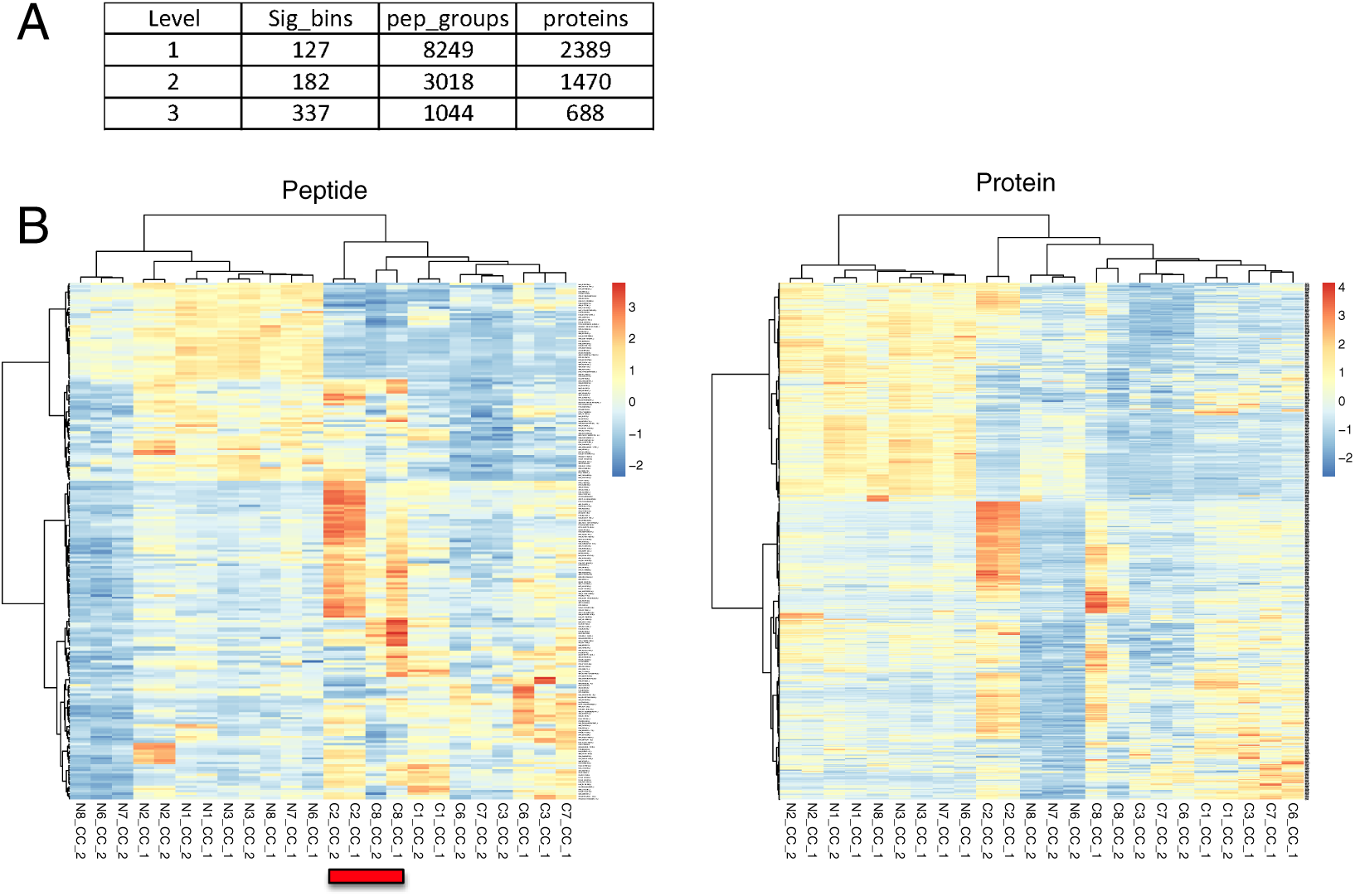
Peptide/protein features associated with MZ-Bin levels. A: Numbers of associated peptides/proteins. With each MZ-Bin iteration, the number of significant features generally increases, with a concomitant decrease in the number of associated peptides/proteins. B: **Class segregation for peptides and proteins**. Hierarchical Clustering (Euclidean distance; Ward’s Linkage) shows that the significant peptides selected based on level-3 MZ-bins can clearly separate the sample classes (Notation: N7_CC_2 refers to normal sample 7, clear-cell renal carcinoma, replicate 2). It is particularly interesting that patients C2 and C8, who suffered from a severe form of the disease, are grouped together. In contrast, the proteins corresponding to these peptides seem to have poorer discrimination, since patients N6, N7 and N8 seem to have been misclassified in the cancer branch. This suggests there is some loss of information in the peptide-to-protein transition.

To investigate whether these peptides, and their associated proteins are informative, we performed hierarchical clustering on the samples. For peptides, the abundance information was based on the SWATH intensity values while for proteins, it was the top two most abundant peptides per sample. We find that for peptides, the separation between sample classes was well-defined. In particular, patients 2 and 8, who suffered from a severe form of renal cancer were grouped together (Figure 4B left). This confirmed that the peptide features associated with the MZ-bins are informative, and can pin-point relevant peptide features.

Interestingly, the class segregation effects are less pronounced for the protein-based clustering. One possibility is that the protein abundance levels for these were not determined based on the significant peptides. If they are based on other peptides and those peptides are insignificant, there are some possibilities: 1/ these other peptides are more likely to have data holes. That is, not consistently measureable across samples. Or 2/, these other peptides are not differentially expressed (so the significant peptides may suggest interesting splice variants).

Critically, the MZ-Bin strategy highlights a problem in proteomics analysis --- In the peptide/protein conundrum, it is expected that without complete protein coverage, it is naïve to quantify proteins based on a limited set of peptide information. Yet, for simplicity of analysis, peptide expression is often ignored in favor of the likely inaccurate protein expression approximations. Here, we showed that the peptide-to-protein transition itself can be a source of quantitation error that can confound analysis.

### Peptide-focused analysis reveals powerful insights into splice variants --- novel splice forms of MAPT associated with good and poor renal cancer outcomes

In the previous section, we found that MZ-Bin associated peptides have stronger signal and thus able to recover the underlying sample classes better. In SWATH, proteins are quantified by the top two peptides with highest intensity in each tissue. However, in different tissues, the top two peptides might not be the same. If they are not the same, perhaps, novel splice forms in each tissue may confound protein quantification.

To identify novel splice variants, we isolated all SWATH-identified peptides that can be unambiguously mapped to the 688 level-3 proteins. For any of these proteins, if all constituent peptides are similarly up or down-regulated, then the abundance of the entire protein is likely to be regulated at the transcriptional level. However, if the constituent peptides are inconsistently expressed, this may be indicative of alternative splice events.

For each peptide, a ratio is calculated based on the division of the median intensity values of the cancer class against the median of the normal class. NAs are substituted by a value of 0.0000001. Each protein is then represented by a series of ratios corresponding to constituent peptide expressions. If the majority of values in either class are dominated by NAs (at least 7/12 in either class), then the median for either class will be extremely small (0.0000001). Thus, generating cancer/normal ratios that will be either extremely large or small. A protein that is characterized by ratios that have extreme values throughout its constituent peptides is unlikely to be reliably quantified, and thus, can be removed from analysis easily. On the other hand, if we simply omitted the NAs during median calculation, it is easy to bias the quantitation on the strength of few observations. We counted the number of NAs for each peptide location and found that the majority of peptides do not have high NA counts; cf. Supplementary Figure 3A. Therefore, for most peptides, we do not expect that excluding NAs will have any strong effects.

It is interesting to note that most of the 688 proteins do not have consistent constituent peptide abundance (Supplementary Data 1). While these proteins are expect to have at least one differential MZ-Bin associated peptide, many of the other constituent peptides are in fact non-differential. To refine the search for relevant alternatively spliced proteins, we introduced the following rules:

1/ there must be at least 10 constituent peptides (to ensure reasonable coverage);
2/ the peptides must be unambiguously mappable to the corresponding protein;
3/ at least 30% constituent peptides are over-expressed, i.e. > 1.25; and
4/ at least 30% constituent peptides are repressed, i.e., < 0.8.

Four proteins met this rule set. These are microtubule-associated protein tau (MAPT, P10636), Heat shock 70 kDa protein 4L (HSPA4L, O95757), Vesicle-fusing ATPase (NSF, P46459) and T-complex protein 1 subunit eta (CCT7, Q99832).

To first understand how the constituent peptides might differentially distinguish normal and cancer classes, we extracted the constituent peptide expression and clustered the patients according to these using Hierarchical clustering; Ward’s linkage and Euclidean distance (Supplementary Figure 3B). However, with the exception of MAPT, the discriminatory power of these peptides for the underlying classes are not particularly strong.

Figure 5A shows that for MAPT, there is differential enrichment of different peptides for severe cancer (C2 and C8 --- highlighted in red) and less severe forms (highlighted in orange). MAPT promotes microtubule assembly and stability, and might be involved in the establishment and maintenance of neuronal polarity. It is commonly associated with neurological diseases such as dementia but its alternative splice forms are already well-described (16). However, despite the relative scarcity of association of MAPT with renal cancer, MAPT has been described as part of a gene signature for predicting severe clear cell renal carcinoma, which is exactly the same renal cancer type our samples were derived from (17). Kosari et al reported MAPT to be down-regulated in aggressive ccRCC (renal cancer). This is consistent with our observation.

**Figure 5.**
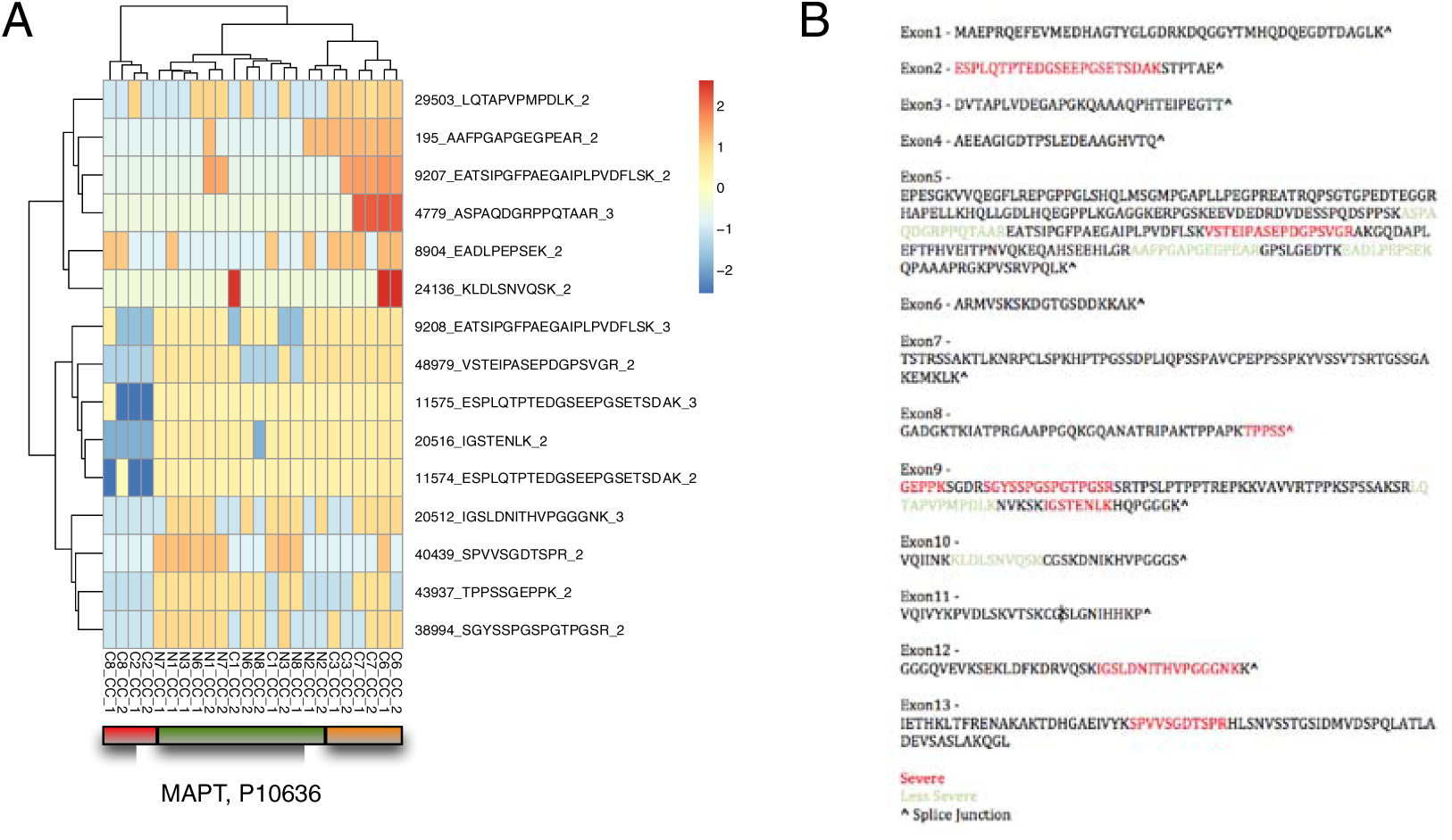
Peptide features associated with MAPT. A: Hierarchical clustering using MAPT peptides. MAPT peptides are differential between severe (red) and less severe cancers (orange). This suggests that these peptides may be useful as markers for prognosis. **B: Localization of MAPT differential peptides within exon junction**. For the most part, severe and less severe peptides are located within different exons, except for exons 5 and 9. This suggests that there may be patients with mutations within these regions that may generate novel splice sites within these exons.

The peptides that are discriminative for severe and less severe forms of renal cancer are evenly distributed along the MAPT protein sequence (Supplementary Figure 4). Hence, we want to know if these discriminative peptides are localized within the splice junctions. To do this, we used genewise to map the MAPT protein sequence against the Mapt unspliced DNA sequence (Supplementary Data 2) where we predicted 13 exons (Figure 5B) (18). For the most part, peptides discriminative for severe and less severe peptides respectively are located on different exons with the exception of exon 5 and exon 9. This may suggest that splicing mechanisms may also be disrupted during tumorigenic progression, or point to the existence of mutations that may generate a novel splice site in some patients within these exons. Certainly, now that we know the peptides that are disrupted, the search space for severe-cancer associated SNPs will be greatly reduced.

We mentioned earlier that MAPT is known to have alternate splice forms. To see if our differential peptides might correspond to any of these splice forms, we picked the two strongest peptides from “severe” and “less severe” groups based on Figure 5 (ASPAQDGRPPQTAAR and KLDLSNVQSK for the less severe group and ESPLQTPTEDGSEEPGSETSDAK and IGSTENLK to represent the severe group). We compared these sequences to eight known splice forms of MAPT (derived from UniprotKb, Supplementary Data 3) using T-coffee multiple sequence aligner (default parameters) (19).

For the peptides associated with severe cancer, ESPLQTPTEDGSEEPGSETSDAK is found on splice forms 4, 5, 7, 8 and 9,while IGSTENLK is found across all splice forms. For the peptides associated with less severe cancer, ASPAQDGRPPQTAAR is found across all splice forms while KLDLSNVQS is found only in splice forms 6,7,8 and 9. While forms 7, 8 and 9 are common to both peptide groups. Forms 4 and 5 are unique to severe cancer associated peptides while form 6 is unique to less severe cancer. Perhaps it is these splice forms themselves that are differentially expressed.

We discuss first ASPAQDGRPPQTAAR (less severe) and IGSTENLK (severe), which are found in all splice forms. These two peptides make perfect biomarkers for the following reasons: 1/ they can be detected in everyone. 2/ their abundance is completely distinct between less severe, severe and normal tissue.

Using Interpro (20, 21) to predict domain information, ASPAQDGRPPQTAAR (less severe) is found on exon 5 (Figure 5B), and corresponds to the Microtubule associated protein MAP2/MAP4/Tau (IPR027324). This domain has a net negative charge and exerts a long-range repulsive force. This provides a mechanism that can regulate microtubule spacing which might facilitate efficient organelle transport (22). IGSTENLK (severe) is found on exon 9 (Figure 5B) and corresponds to several domains --- IPR027324 (as seen earlier), Microtubule-associated protein Tau (IPR002955) which is involved in microtubule binding, and Microtubule associated protein, tubulin-binding repeat (IPR001084) which is implicated in tubulin binding and which seem to have a stiffening effect on microtubules.

Exon 5 has a propensity for peptides associated with less severe cancer (3), which implies its over-expression is associated with less severe cancer. Disrupting this, as detected with VSTEIPASEPDGPSVGR, could potentially reverse this either by mutating part of the exon, or by overall down-regulation of this region.

Exon 9 on the other hand, has a propensity for peptides associated with severe cancer (3 as well), which implies its repression is needed for severe cancer to progress. Similarly, a single peptide, TAPVPMPDLK is found to be overexpressed and associated with less severe phenotype. The repeat domain (IPR001084, Microtubule associated protein, tubulin-binding repeat) is usually located on the C-terminus, repeated in tandem about 3 to 4 times, and is implicated in tubulin binding and seem to have a stiffening effect on microtubules. Perhaps, over-expression of IPR001084 stabilizes the microtubules, and makes it harder for the cancer to undergo metastasis.

To discover if there are any more interesting domain-specific information associated with the alternative splice forms, we aligned the sequences of splice forms 4, 5 (severe) and 6 (less severe) using t-coffee. We then extracted two representative domain sequences --- ESPLQTPTEDGSEEPGSETSDAKSTPTAEDVTAPLVDEGAPGKQAAAQPHTEIPEGTT for severe (severe domain) and QIINKKLDLSNVQSKCGSKDNIKHVPGGGSV for less severe (less severe domain). As before, we checked if these corresponded to any known protein domains or annotated functional sequences using Interpro (20, 21).

The severe domain is found on exon 2 and corresponded to IPR027324 while the less severe domain is found on exon 10 and maps to both IPR027324 and IPR001084. As before, the repression of exon 2 suggests this region is potentially impaired in severe cancer. In a similar vein, overexpression of exon 10 may increase microtubule stability, making it difficult for cells to effect metastasis.

In the literature, upregulated MAPT is a good-prognosis indicator in renal cancer while its down-regulation might mean the opposite (23). The altered regulation is in fact not consistent across its entire length. Moreover, specific peptide regions are over-expressed for less severe renal cancer while other non-overlapping regions are repressed for severe renal cancer. Consideration of these specific peptide regions is more useful as prognostic markers than using the entire protein length, which may dilute the signal.

### Differential MZ-Bin associated proteins are more likely to be true features

We’ve shown earlier that MZ-Bin associated peptides have stronger class-differential power, and that these peptides can be used for identifying useful splice forms of the corresponding proteins. On the other hand, corresponding proteins have weaker signal (688 proteins), possibly due to the naïve belief that constituent peptides are a good approximation for total protein expression.

Earlier, we also noted that deploying simple t-test (SP) on the proteins alone yielded an unusually large number of significant features (1,649 out of 3,123 proteins). About 1,247 proteins are SP-unique, and not found in the set of MZ-Bin associated proteins.

In earlier work, we’ve shown that networks are useful for removing unreliable proteins since proteins that do work together tends to belong to the same subnetworks (24–28). We’ve also shown that real biological complexes are far superior to predicted subnetworks (29).

Using 1,363 protein complexes derived from CORUM (30), a vector of intersections was generated for both the SP-unique and MZ-Bin lists. These intersection vectors were summed and divided over the complexes in which an intersection of at least 1 was observed (The less complexes the proteins are distributed over, the greater the value). This is then normalized by the number of proteins in the SP-unique and MZ-Bin lists respectively to generate Sum_hit-rate_SP-unique_ and Sum_hit-rate_MZ-Bin_ respectively. The ratio of Sum_hit-rate_MZ-Bin_ over Sum_hit-rate_SP-unique_ was 1.4x, indicating that there is stronger enrichment for same complexes in MZ-Bin proteins over those unique to the protein-based feature selection. Since proteins tend to work together in groups, this suggests that the proteins identified by MZ-Bin are higher quality.

## Conclusions and future work

MZ-Bin is a practical approach towards feature selection in proteomic experiments. Significant MZ-bins are informative, and can be tracked to peptides/proteins relevant to the phenotype.

We’ve also demonstrated that peptides have stronger signals than their respective proteins, and revealed some potential flaws in the standard peptide-to-protein quantitation approach.

The MZ-Bin approach allows the identification of relevant proteins. Checking the expression values of the constituent peptides allows determination of potential splice variants. We demonstrate the checking for splice variants is rewarding, and uncovered for the first time, how splice variants are implicated in severe and less severe renal cancer.

MZ-Bin is still in its infancy. We believe that the inverted approach towards proteomics analysis has yet to achieve its full potential. In future work, the following advances are needed:

1/ Develop more powerful feature-identification algorithms with high statistical power and sensitivity to ensure all bins containing useful signal are retained, and not lost. And develop a means of checking the reproducibility of the identified feature sets at each bin level.
2/ Develop more complete libraries for comprehensive mapping of spectral features.
3/ Consider how MS2 spectra can be co-deployed in this analysis strategy.
4/ Consider how PTM analysis can be incorporated into this framework.
5/ Consider how to incorporate de novo sequencing techniques using the unannotated MZ-bins as “seed points”.

## Acknowledgements

WWBG is funded by a Professorship of Bioinformatics, School of Pharmaceutical Science and Technology, Tianjin University, China. LW is funded in part by a Singapore Ministry of Education tier-2 grant, MOE2012-T2-1-061.

## Author contributions

WWBG co-designed and implemented the MZ-Bin pipeline, performed analysis, and wrote the manuscript. LW co-designed MZ-Bin, co-wrote the manuscript.

## Supplementary figures

**Supplementary Figure 1.**
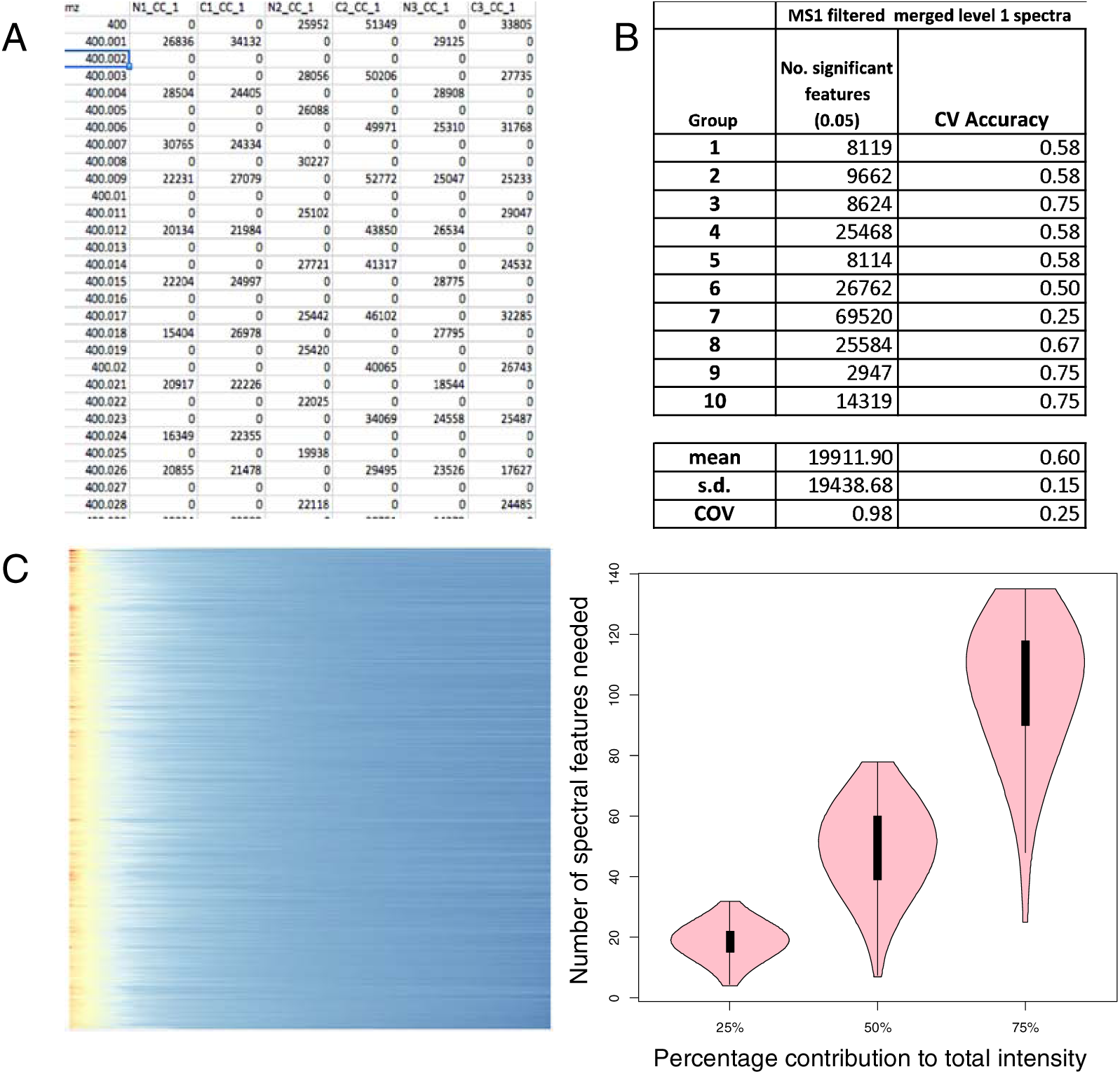
A: Raw spectra has high data holes. Due to stochastic effects and misalignments, many of the m/z values have high presence of data holes (0s). **B: Unbinned features are poorly predictive.** Without binning, many of the features are misaligned, generating many data holes and therefore poorly predictive. The cross-validation accuracy as shown here is about 0.6. This is low. **C: Contribution of signal intensities**. The heatmap on the left shows that the majority of signal per MZ-Bin is contributed by a small number of features. The violinplot on the right shows the distribution of spectral features required to exceed 25%, 50% and 75% of each MZ-bin’s total intensity. This confirms that a small number of features dominates the signal in each MZ-bin.

**Supplementary Figure 2.**
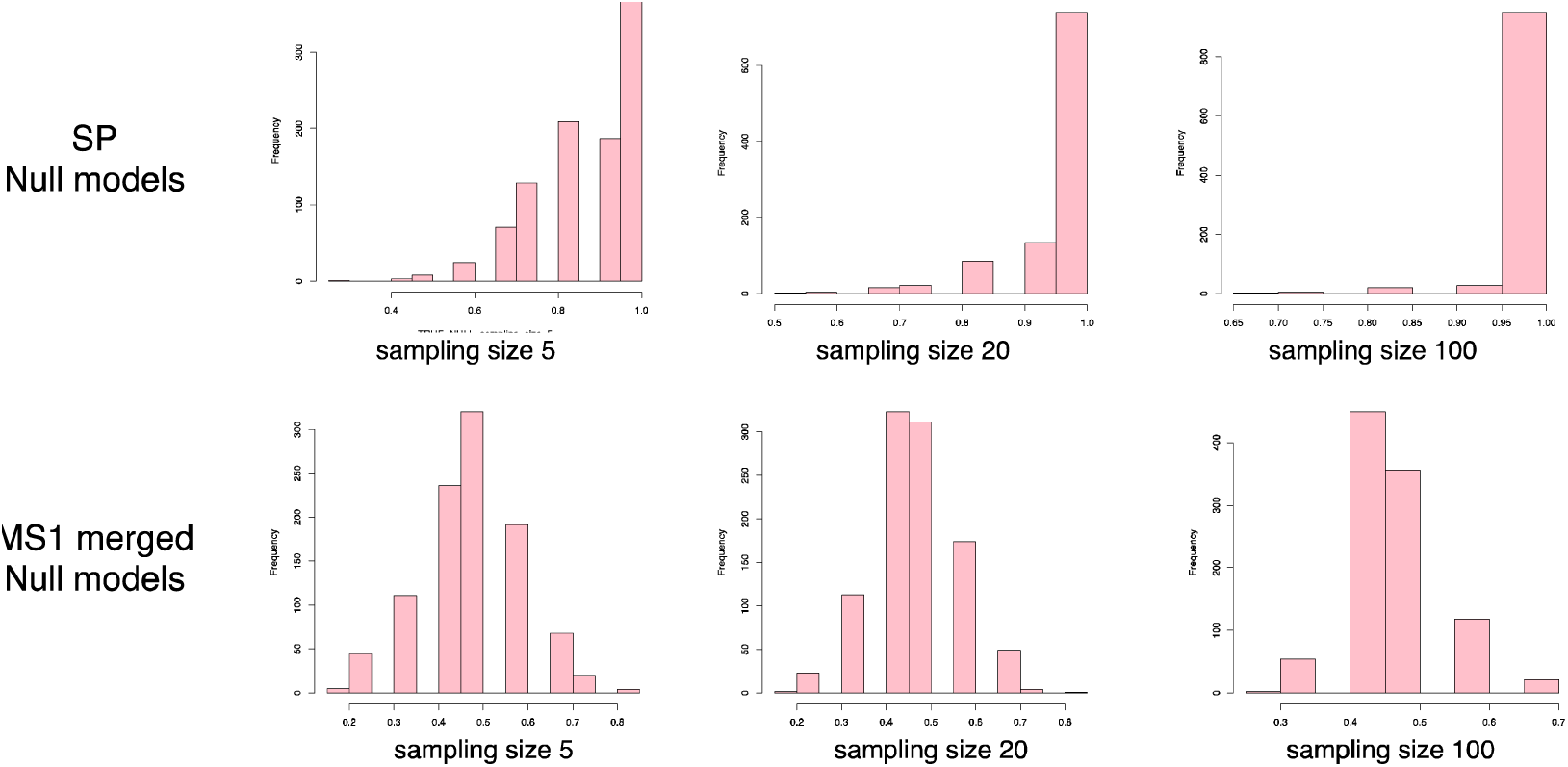
Null models for cross-validation prediction accuracy. In SP, any random selection of at least 5 proteins can build a model with very high cross-validation accuracy, i.e., any random selection of proteins are highly class-predictive. In contrast, most MZ-bins (MS1-merged level 1 is shown here) are not correlated with the classes and, thus, randomization generates more evenly distributed null CV values.

**Supplementary Figure 3.**
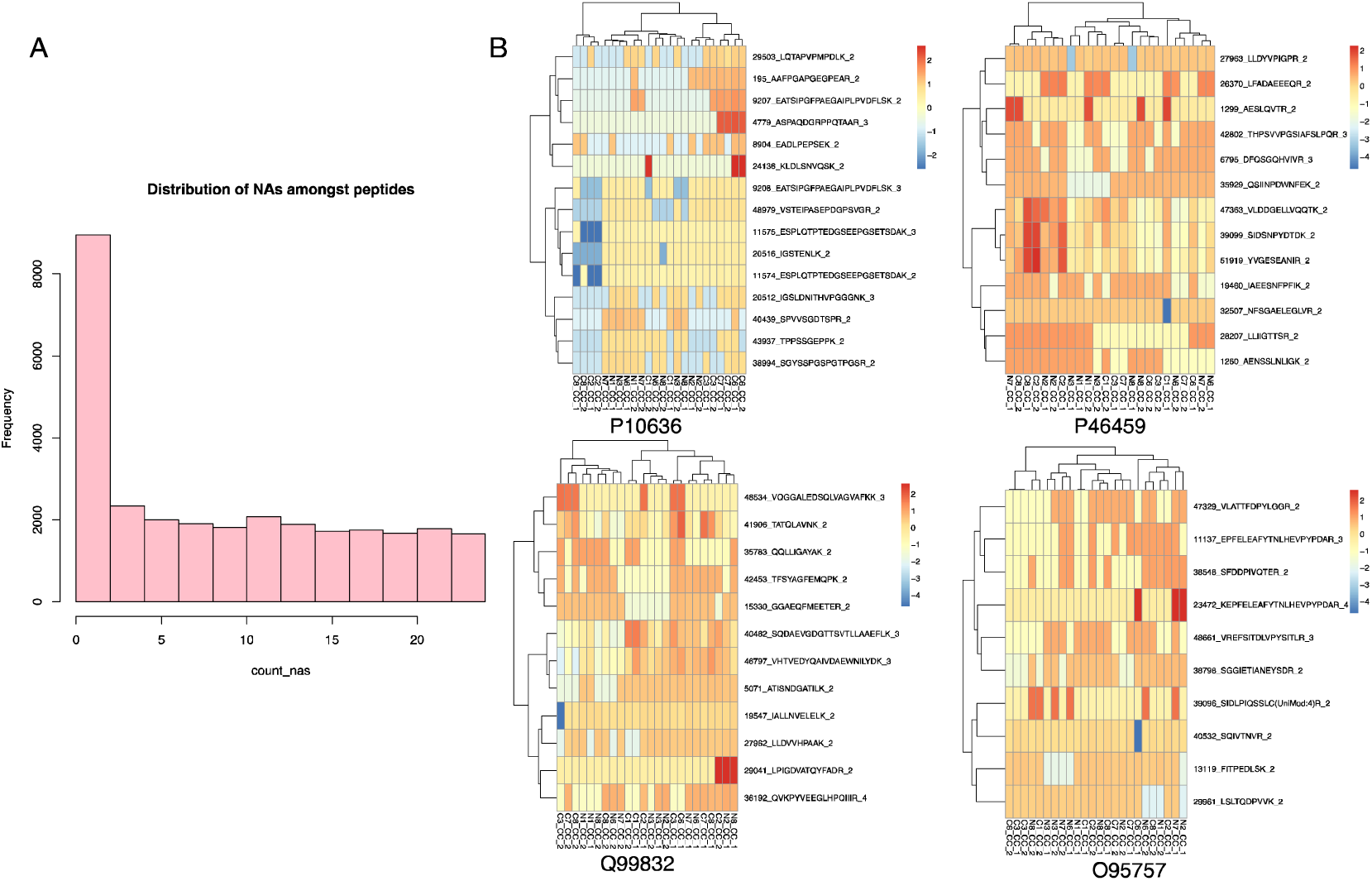
A: NA counts for each peptide group. Most peptides are well-quantitiated across samples. Hence we do not expect that excluding NAs from median calculations will have a strong effect**. B. Hierarchical clustering for 4 altnerative splice candidates.** Although these 4 proteins met our filtering criteria for differentially expressed constituent peptides, only the first (P10636) can discriminate between normal, severe and less severe cancer classes.

**Supplementary Figure 4.**
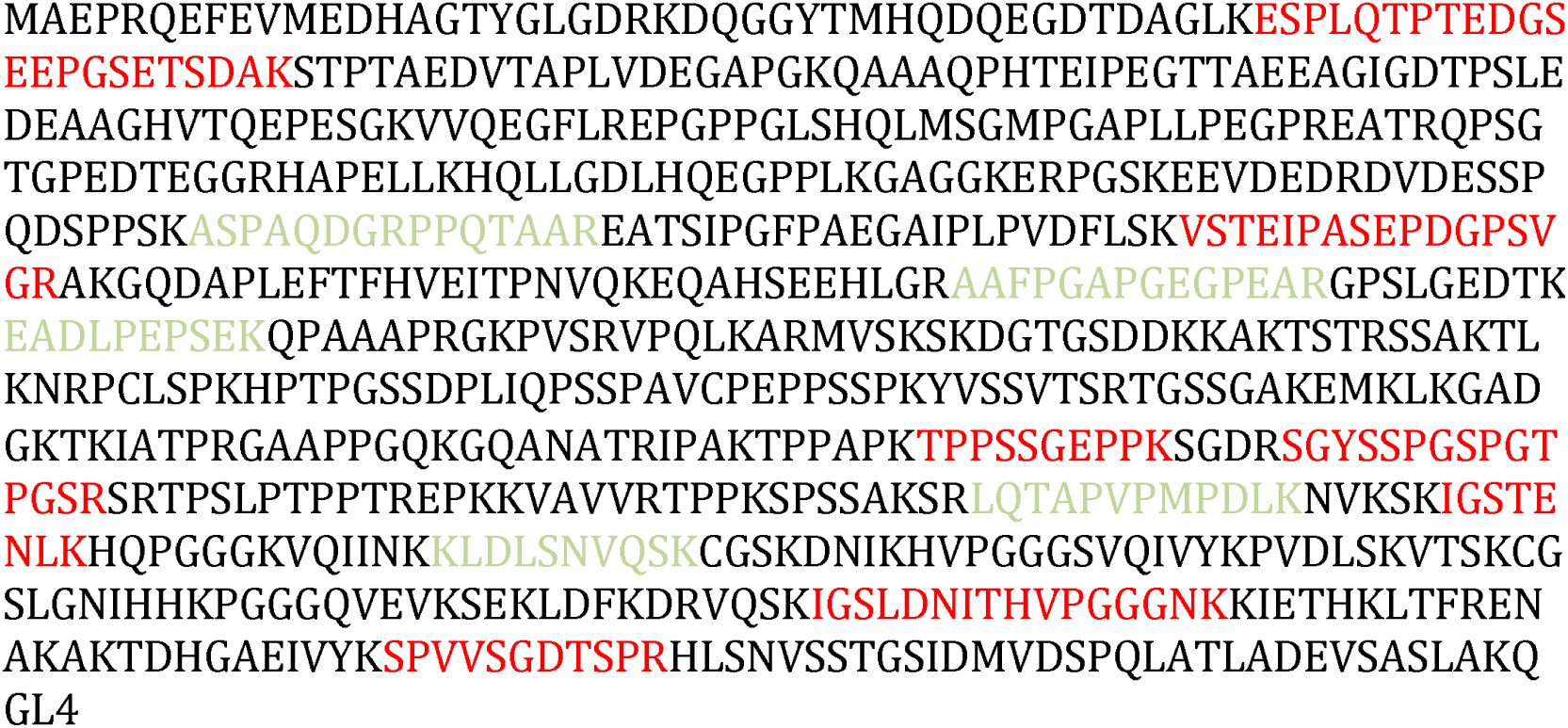
Protein sequence for MAPT. The distribution of severe and less severe peptides (highlighted in red and green respectively) are quite well spaced. We hypothesize that these peptides are interspersed within different splice sites (exons).

## Supplementary tables

**Supplementary Table 1.** Combined tables for both level-1 MS1 and MS2, merged and unmerged at 95% and 99% significance levels.

| Group | MS1 merged level 1 spectra |  |  | MS1 merged level 1 spectra |  |  | MS2 merged level 1 spectra |  |  | MS2 merged level 1 spectra |  |  |
| --- | --- | --- | --- | --- | --- | --- | --- | --- | --- | --- | --- | --- |
|  | No. significant features (0.05) | self_validation | cross_validation | No. significant features (0.01) | self_validation | cross_validation | No. significant features (0.05) | self_validation | cross_validation | No. significant features (0.01) | self_validation | cross_validation |
| 1 | 103 | 0.92 | 0.83 | 20 | 0.92 | 0.83 | 167 | 1.00 | 0.83 | 83.00 | 1.00 | 0.83 |
| 2 | 228 | 1.00 | 0.83 | 57 | 1.00 | 0.75 | 177 | 1.00 | 0.83 | 59.00 | 1.00 | 0.83 |
| 3 | 397 | 0.92 | 0.75 | 62 | 0.92 | 0.83 | 1328 | 0.92 | 0.83 | 581.00 | 1.00 | 0.83 |
| 4 | 757 | 1.00 | 0.75 | 400 | 1.00 | 0.75 | 1428 | 0.92 | 0.83 | 755.00 | 1.00 | 0.83 |
| 5 | 376 | 1.00 | 0.75 | 198 | 1.00 | 0.75 | 336 | 1.00 | 0.83 | 119.00 | 1.00 | 0.83 |
| 6 | 237 | 1.00 | 0.83 | 132 | 1.00 | 0.75 | 100 | 1.00 | 0.83 | 51.00 | 1.00 | 0.83 |
| 7 | 137 | 0.92 | 0.92 | 38 | 0.92 | 0.92 | 68 | 0.92 | 0.83 | 26.00 | 0.92 | 0.83 |
| 8 | 248 | 1.00 | 0.83 | 79 | 1.00 | 0.83 | 121 | 0.92 | 0.83 | 36.00 | 0.92 | 0.83 |
| 9 | 654 | 1.00 | 0.75 | 451 | 1.00 | 0.75 | 1354 | 1.00 | 0.83 | 961.00 | 1.00 | 0.83 |
| 10 | 321 | 1.00 | 0.83 | 71 | 1.00 | 0.83 | 855 | 1.00 | 0.83 | 295.00 | 1.00 | 0.83 |
| mean | 345.80 |  | 0.81 | 150.80 |  | 0.80 | 593.40 |  | 0.83 | 296.60 |  | 0.83 |
| s.d. | 212.41 |  |  | 153.78 |  |  | 581.88 |  |  | 344.40 |  |  |
| COV | 0.61 |  |  | 1.02 |  |  | 0.98 |  |  | 1.16 |  |  |

| Group | MS1 unmerged level 1 spectra |  |  | MS1 unmerged level 1 spectra |  |  | MS2 unmerged level 1 spectra |  |  | MS2 unmerged level 1 spectra |  |  |
| --- | --- | --- | --- | --- | --- | --- | --- | --- | --- | --- | --- | --- |
|  | No. significant features (0.05) | self_validation | cross_validation | No. significant features (0.01) | self_validation | cross_validation | No. significant features (0.05) | self_validation | cross_validation | No. significant features (0.01) | self_validation | cross_validation |
| 1 | 95 | 0.92 | 0.83 | 20 | 0.92 | 0.83 | 4667 | 0.92 | 0.92 | 955 | 0.92 | 0.83 |
| 2 | 178 | 1.00 | 0.83 | 50 | 1.00 | 0.75 | 11605 | 1.00 | 0.83 | 4743 | 1.00 | 0.83 |
| 3 | 326 | 0.92 | 0.75 | 57 | 0.92 | 0.83 | 17501 | 0.83 | 0.75 | 3155 | 0.92 | 0.83 |
| 4 | 567 | 1.00 | 0.75 | 310 | 1.00 | 0.75 | 30378 | 1.00 | 0.75 | 14539 | 1.00 | 0.75 |
| 5 | 302 | 1.00 | 0.75 | 118 | 1.00 | 0.75 | 15036 | 1.00 | 0.75 | 7678 | 1.00 | 0.75 |
| 6 | 203.00 | 1.00 | 0.83 | 132 | 1.00 | 0.75 | 6595 | 1.00 | 0.83 | 2953 | 1.00 | 0.75 |
| 7 | 120 | 0.92 | 0.92 | 36 | 0.92 | 0.92 | 3806 | 1.00 | 0.92 | 1356 | 1.00 | 0.92 |
| 8 | 208 | 1.00 | 0.83 | 74 | 1.00 | 0.83 | 5690 | 1.00 | 0.83 | 1861 | 1.00 | 0.83 |
| 9 | 532 | 1.00 | 0.75 | 381 | 1.00 | 0.75 | 29052 | 1.00 | 0.67 | 15366 | 1.00 | 0.67 |
| 10 | 245 | 1.00 | 0.83 | 51 | 1.00 | 0.83 | 13458 | 1.00 | 0.92 | 3664 | 1.00 | 0.83 |
| mean | 277.6 |  | 0.81 | 122.9 |  | 0.80 | 13778.80 |  | 0.82 | 5627.00 |  | 0.80 |
| s.d. | 160.202095 |  |  | 123.417494 |  |  | 9595.05 |  |  | 5277.06 |  |  |
| COV | 0.57709688 |  |  | 1.00421069 |  |  | 0.70 |  |  | 0.94 |  |  |

**Supplementary Table 2.** Comparison of level-1 MZ-bins following removal of low-quality SWATH windows in MS1 space. Following the removal of low-quality SWATH windows (cf. Figure 3), the numbers of selected features for both SWATH window-merged and un-merged level-1 MZ-bins now match exactly.

|  | MS1 filtered unmerged level 1 spectra |  |  | MS1 filtered merged level 1 spectra |  |  |
| --- | --- | --- | --- | --- | --- | --- |
| Group | No. significant features (0.05) | CV Accuracy | CV p-value | CV Accuracy | CV Accuracy | CV p-value |
| 1 | 82.00 | 0.83 | 0.00 | 82.00 | 0.83 | 0.00 |
| 2 | 130.00 | 0.75 | 0.00 | 130.00 | 0.75 | 0.00 |
| 3 | 179.00 | 0.75 | 0.00 | 179.00 | 0.75 | 0.00 |
| 4 | 234.00 | 0.75 | 0.00 | 234.00 | 0.75 | 0.00 |
| 5 | 220.00 | 0.75 | 0.00 | 220.00 | 0.75 | 0.00 |
| 6 | 158.00 | 0.75 | 0.00 | 158.00 | 0.75 | 0.00 |
| 7 | 96.00 | 0.92 | 0.00 | 96.00 | 0.92 | 0.00 |
| 8 | 166.00 | 0.83 | 0.00 | 166.00 | 0.83 | 0.00 |
| 9 | 230.00 | 0.75 | 0.00 | 230.00 | 0.75 | 0.00 |
| 10 | 98.00 | 0.92 | 0.00 | 98.00 | 0.92 | 0.00 |
| mean | 159.30 | 0.80 | 0.00 | 159.30 | 0.80 | 0.00 |
| s.d. | 167.89 | 0.80 | 0.00 | 167.89 | 0.80 | 0.00 |
| COV | 171.14 | 0.80 | 0.00 | 171.14 | 0.80 | 0.00 |

## References

1. Sajic, T.; Liu, Y.; Aebersold, R., Using data-independent, high-resolution mass spectrometry in protein biomarker research: perspectives and clinical applications. Proteomics Clin Appl 2015, 9, (3–4), 307–21.

2. Goh, W. W.; Wong, L., Networks in proteomics analysis of cancer. Curr Opin Biotechnol 2013, 24, (6), 1122–8.

3. Goh, W. W.; Wong, L., Computational proteomics: designing a comprehensive analytical strategy. Drug Discov Today 2014, 19, (3), 266–74.

4. Goh, W. W.; Wong, L.; Sng, J. C., Contemporary network proteomics and its requirements. Biology (Basel) 2013, 3, (1), 22–38.

5. Saeed, F.; Hoffert, J. D.; Knepper, M. A., CAMS-RS: Clustering Algorithm for Large-Scale Mass Spectrometry Data using Restricted Search Space and Intelligent Random Sampling. IEEE/ACM Trans Comput Biol Bioinform 2013.

6. Sirota, F. L.; Batagov, A.; Schneider, G.; Eisenhaber, B.; Eisenhaber, F.; Maurer-Stroh, S., Beware of moving targets: reference proteome content fluctuates substantially over the years. J Bioinform Comput Biol 2012, 10, (6), 1250020.

7. Nesvizhskii, A. I.; Vitek, O.; Aebersold, R., Analysis and validation of proteomic data generated by tandem mass spectrometry. Nat Methods 2007, 4, (0), 787–97.

8. Saeed, F.; Hoffert, J. D.; Pisitkun, T.; Knepper, M. A., Exploiting Thread-Level and Instruction-Level Parallelism to Cluster Mass Spectrometry Data using Multicore Architectures. Netw Model Anal Health Inform Bioinform 2014, 3, 54. 9.

9. Plumb, R. S.; Johnson, K. A.; Rainville, P.; Smith, B. W.; Wilson, I. D.; Castro-Perez, J. M.; Nicholson, J. K., UPLC/MS(E); a new approach for generating molecular fragment information for biomarker structure elucidation. Rapid Commun Mass Spectrom 2006, 20, (3), 1989–94.

10. Gillet, L. C.; Navarro, P.; Tate, S.; Rost, H.; Selevsek, N.; Reiter, L.; Bonner, R.; Aebersold, R., Targeted data extraction of the MS/MS spectra generated by data-independent acquisition: a new concept for consistent and accurate proteome analysis. Mol Cell Proteomics 2012, 11, (6), O111 016717.

11. Guo, T.; Kouvonen, P.; Koh, C. C.; Gillet, L. C.; Wolski, W. E.; Rost, H. L.; Rosenberger, G.; Collins, B. C.; Blum, L. C.; Gillessen, S.; Joerger, M.; Jochum, W.; Aebersold, R., Rapid mass spectrometric conversion of tissue biopsy samples into permanent quantitative digital proteome maps. Nat Med 2015.

12. Collins, B. C.; Gillet, L. C.; Rosenberger, G.; Rost, H. L.; Vichalkovski, A.; Gstaiger, M.; Aebersold, R., Quantifying protein interaction dynamics by SWATH mass spectrometry: application to the 14-3-3 system. Nat Methods 2013, 10, (2), 1246–53.

13. Rost, H. L.; Rosenberger, G.; Navarro, P.; Gillet, L.; Miladinovic, S. M.; Schubert, O. T.; Wolski, W.; Collins, B. C.; Malmstrom, J.; Malmstrom, L.; Aebersold, R., OpenSWATH enables automated, targeted analysis of data-independent acquisition MS data. Nat Biotechnol 2014, 32, (3), 219–23.

14. Guo, T.; Kouvonen, P.; Koh, C. C.; Gillet, L. C.; Wolski, W. E.; Rost, H. L.; Rosenberger, G.; Collins, B. C.; Blum, L. C.; Gillessen, S.; Joerger, M.; Jochum, W.; Aebersold, R., Rapid mass spectrometric conversion of tissue biopsy samples into permanent quantitative digital proteome maps. Nat Med 2015, 21, (4), 407–13.

15. Raju, T. N., William Sealy Gosset and William A. Silverman: two “students” of science. Pediatrics 2005, 116, (3), 732–5.

16. Garcia-Blanco, M. A.; Baraniak, A. P.; Lasda, E. L., Alternative splicing in disease and therapy. Nat Biotechnol 2004, 22, (5), 535–46.

17. Kosari, F.; Parker, A. S.; Kube, D. M.; Lohse, C. M.; Leibovich, B. C.; Blute, M. L.; Cheville, J. C.; Vasmatzis, G., Clear cell renal cell carcinoma: gene expression analyses identify a potential signature for tumor aggressiveness. Clin Cancer Res 2005, 11, (4), 5128–39.

18. Birney, E.; Clamp, M.; Durbin, R., GeneWise and Genomewise. Genome Res 2004, 14, (5), 988–95.

19. Notredame, C.; Higgins, D. G.; Heringa, J., T-Coffee: A novel method for fast and accurate multiple sequence alignment. J Mol Biol 2000, 302, (1), 205–17.

20. Mitchell, A.; Chang, H. Y.; Daugherty, L.; Fraser, M.; Hunter, S.; Lopez, R.; McAnulla, C.; McMenamin, C.; Nuka, G.; Pesseat, S.; Sangrador-Vegas, A.; Scheremetjew, M.; Rato, C.; Yong, S. Y.; Bateman, A.; Punta, M.; Attwood, T. K.; Sigrist, C. J.; Redaschi, N.; Rivoire, C.; Xenarios, I.; Kahn, D.; Guyot, D.; Bork, P.; Letunic, I.; Gough, J.; Oates, M.; Haft, D.; Huang, H.; Natale, D. A.; Wu, C. H.; Orengo, C.; Sillitoe, I.; Mi, H.; Thomas, P. D.; Finn, R. D., The InterPro protein families database: the classification resource after 15 years. Nucleic Acids Res 2015, 43, (Database issue), D213–21.

21. Apweiler, R.; Attwood, T. K.; Bairoch, A.; Bateman, A.; Birney, E.; Biswas, M.; Bucher, P.; Cerutti, L.; Corpet, F.; Croning, M. D.; Durbin, R.; Falquet, L.; Fleischmann, W.; Gouzy, J.; Hermjakob, H.; Hulo, N.; Jonassen, I.; Kahn, D.; Kanapin, A.; Karavidopoulou, Y.; Lopez, R.; Marx, B.; Mulder, N. J.; Oinn, T. M.; Pagni, M.; Servant, F.; Sigrist, C. J.; Zdobnov, E. M., The InterPro database, an integrated documentation resource for protein families, domains and functional sites. Nucleic Acids Res 2001, 29, (1), 37–40.

22. Mukhopadhyay, R.; Hoh, J. H., AFM force measurements on microtubule-associated proteins: the projection domain exerts a long-range repulsive force. FEBS Lett 2001, 505, (3), 374–8.

23. Brooks, S. A.; Brannon, A. R.; Parker, J. S.; Fisher, J. C.; Sen, O.; Kattan, M. W.; Hakimi, A. A.; Hsieh, J. J.; Choueiri, T. K.; Tamboli, P.; Maranchie, J. K.; Hinds, P.; Miller, C. R.; Nielsen, M. E.; Rathmell, W. K., ClearCode34: A prognostic risk predictor for localized clear cell renal cell carcinoma. Eur Urol 2014, 66, (1), 77–84.

24. Goh, W. W.; Lee, Y. H.; Zubaidah, R. M.; Jin, J.; Dong, D.; Lin, Q.; Chung, M. C.; Wong, L., Network-based pipeline for analyzing MS data: an application toward liver cancer. J Proteome Res 2011, 10, (5), 2261–72.

25. Goh, W. W.; Lee, Y. H.; Ramdzan, Z. M.; Sergot, M. J.; Chung, M.; Wong, L., Proteomics signature profiling (PSP): a novel contextualization approach for cancer proteomics. J Proteome Res 2012, 11, (3), 1571–81.

26. Goh, W. W.; Lee, Y. H.; Ramdzan, Z. M.; Chung, M. C.; Wong, L.; Sergot, M. J., A network-based maximum link approach towards MS identifies potentially important roles for undetected ARRB1/2 and ACTB in liver cancer progression. Int J Bioinform Res Appl 2012, 8, (3), 155–70.

27. Goh, W. W.; Lee, Y. H.; Chung, M.; Wong, L., How advancement in biological network analysis methods empowers proteomics. Proteomics 2012, 12, (4–5), 550–63.

28. Goh, W. W.; Fan, M.; Low, H. S.; Sergot, M.; Wong, L., Enhancing the utility of Proteomics Signature Profiling (PSP) with Pathway Derived Subnets (PDSs), performance analysis and specialised ontologies. BMC Genomics 2013, 14, 35. 29.

29. Goh, W. W.; Sergot, M. J.; Sng, J. C.; Wong, L., Comparative network-based recovery analysis and proteomic profiling of neurological changes in valproic Acid-treated mice. J Proteome Res 2013, 12, (5), 2116-27.

30. Ruepp, A.; Waegele, B.; Lechner, M.; Brauner, B.; Dunger-Kaltenbach, I.; Fobo, G.; Frishman, G.; Montrone, C.; Mewes, H. W., CORUM: the comprehensive resource of mammalian protein complexes--2009. Nucleic Acids Res 2010, 38, (Database issue), D497–501.

